# Work and oxygen consumption of isolated right ventricular papillary muscle in experimental pulmonary hypertension

**DOI:** 10.1101/2021.12.30.474521

**Authors:** Willem J. van der Laarse, Sylvia J.P. Bogaards, Ingrid Schalij, Anton Vonk Noordegraaf, Frédéric M. Vaz, Duncan van Groen

**Affiliations:** Department of Physiology, Amsterdam University Medical Centers,The Netherlands; Department of Pulmonology, Amsterdam University Medical Centers,The Netherlands; Amsterdam Cardiovascular Sciences, VU University Amsterdam, Amsterdam University Medical Centers, Amsterdam, The Netherlands and Laboratory Genetic Metabolic Diseases, Amsterdam UMC, University of Amsterdam, Amsterdam University Medical Centers,The Netherlands; Department of Clinical Chemistry, Amsterdam Gastroentrology Endocrinology Metabolism, Amsterdam, Department of Pediatrics, Amsterdam University Medical Centers,The Netherlands; Emma Children’s Hospital, Amsterdam University Medical Centers, Core Facility Metabolomics, Amsterdam University Medical Centers,The Netherlands

**Keywords:** Pulmonary hypertension, monocrotaline, myocardial efficiency, right-sided heart failure, papillary muscle, blebbistatin, oxygen consumption, work loop, mitochondria, cardiolipin, cytochrome c release, cardiolipin

## Abstract

Right-sided myocardial mechanical efficiency (work output/metabolic energy input) in pulmonary hypertension can be severely reduced. We determined the contribution of intrinsic myocardial determinants of efficiency using papillary muscle preparations from monocrotaline-induced pulmonary hypertensive (MCT-PH) rats. The hypothesis was tested that efficiency is reduced by mitochondrial dysfunction in addition to increased activation heat reported previously. Right ventricular (RV) muscle preparations were subjected to 5 Hz sinusoidal length changes at 37°C. Work and suprabasal oxygen consumption (VO_2_) were measured before and after cross-bridge inhibition by blebbistatin. Cytosolic cytochrome c concentration, myocyte cross-sectional area, proton permeability of the inner mitochondrial membrane (PP IMM), and monoamine oxidase (MAO)-A and glucose 6-phosphate dehydrogenase (G-6-PDH) activities and phosphatidylglycerol (PG) and cardiolipin (CL) contents were determined. Mechanical efficiency ranged from 23 to 11% in control (n = 6) and from 22 to 1% in MCT-PH (n = 15) and correlated with work (r^2^ = 0.68, P < 0.0001) but not with VO_2_ (r^2^ = 0.004, P = 0.7919). VO_2_ for cross-bridge cycling was proportional to work (r^2^ = 0.56, P = 0.0005). Blebbistatin-resistant VO_2_ (r^2^ = 0.32, P = 0.0167) and IMM PP (r^2^ = 0.36, P = 0.0110) correlated inversely with efficiency. Together, these variables explained the variance of efficiency (coefficient of multiple determination R^2^ = 0.79, P = 0.0001). Cytosolic cytochrome c correlated inversely with work (r^2^ = 0.28, P = 0.0391), but not with efficiency (r^2^ = 0.20, P = 0.0867). G-6-PDH, MAO-A and PG/CL increased in the RV wall of MCT-PH but did not correlate with efficiency. Reduced myocardial efficiency in MCT-PH is due to activation processes and mitochondrial dysfunction. The variance of work and the ratio of activation heat reported previously and blebbistatin-resistant VO_2_ are discussed.

**Key points:**

- Mechanical efficiency of right ventricular myocardium is reduced in pulmonary hypertension. Increased energy use for activation processes has been demonstrated previously, but the contribution of mitochondrial dysfunction is unknown.
- Work and oxygen consumption is determined during work loops. Oxygen consumption for activation and cross-bridge cycling confirm the previous heat measurements.
- Cytosolic cytochrome c concentration, proton permeability of the mitochondrial inner membrane and phosphatidylglycerol/cardiolipin are increased in experimental pulmonary hypertension.
- Mitochondrial dysfunction in right ventricular myocytes is related to reduced work and mechanical efficiency in experimental pulmonary hypertension.
- Upregulation of the pentose phosphate pathway and a potential gap in the energy balance suggest mitochondrial dysfunction in right ventricular overload is due to excessive production of reactive oxygen species.

## Introduction

Right-sided heart failure is the major cause of death of patients with pulmonary arterial hypertension. Despite progress in diagnosis and treatment, the mechanism of the transition of pulmonary hypertension-induced hypertrophy to failure of the right-sided myocardium remains enigmatic (for review, see Westerhof et al. 2017; Thenappan et al. 2018). The transition to failure in patients coincides with a reduction of right-sided myocardial efficiency calculated as pump work/metabolic energy input (Wong et al. 2011a; Yoshinaga et al. 2014; Ahmadi et al. 2020). Reduced myocardial efficiency can be due to suboptimal right ventricular to pulmonary artery coupling (Vonk Noordegraaf et al. 2017) and/or an intrinsic efficiency reduction as has been demonstrated in experimental models (Cooper et al. 1973; Wong et al. 2010; Pham et al. 2018; Noly et al. 2019).

Reduced efficiency is important because it increases myocardial oxygen demand. The increase of mean pulmonary arterial pressure from 15 to > 70 mmHg in patients doubles right-sided myocardial mass, which can be explained by the increase in cardiomyocyte cross-sectional area (Beltrami et al. 1994, Ruiter et al. 2012). Because the cardiomyocytes do not proliferate to a significant extent, the increased RV afterload and reduced efficiency can increase suprabasal oxygen demand per myocyte more than eightfold. It is unlikely that coronary flow reserve can meet oxygen requirements (Wong et al, 2011b; Ruiter et al. 2012; Sree Raman et al. 2021), implying that maximum cardiac output in patients is limited by oxygen supply.

Oxidative metabolism can limit ATP production either by hypoxia (Oknińska et al. 2021) or otherwise (Balestra et al. 2015; Fowler et al. 2019; Koop et al. 2019). It has been demonstrated that hypoxia-inducible transcription factor (HIF)-1α is upregulated in hypertrophied right ventricular cardiomyocytes (des Tombe et al. 2002; Simonides et al. 2008; Sutendra et al. 2013), complex I of the electron transport chain is inhibited (Wüst et al. 2016) and complex II becomes a source of reactive oxygen species (Redout et al. 2007). Cytochrome c enters the cytosol in cardiomyocytes with cross-sectional area larger than twice control (van Beek-Harmsen & van der Laarse, 2005). The release of cytochrome c is detectable prior to the transition to heart failure, at two weeks after the MCT injection (van Beek-Harmsen et al. 2011). In addition, the function of the phosphocreatine shuttle may be impaired (Ishikawa et al. 1995; Lamberts et al. 2007; Fowler et al. 2015). In MCT-PH models surviving for four weeks or longer, mitochondria in the right-sided myocardium are enlarged (Kajihara, 1970), mitochondrial efficiency (ATP/O_2_) decreases (Raczniak et al, 1977), phosphocreatine and ATP contents are reduced (Daicho et al., 2009), and xanthine oxidase is upregulated (de Jong et al., 2000).

Uncovering early mechanisms reducing efficiency of failing right-sided myocardium is a requirement to develop targeted therapy. Ruthenium red has been used by Cooper et al. (1973) to demonstrate in isolated mitochondria that futile calcium cycling may be causing increased oxygen consumption in overloaded right-sided myocardium of the cat. Pham et al. (2018) have shown that activation heat of rat myocardial trabeculae from the right ventricle is increased in MCT-PH, suggesting increased Ca^2+^ cycling for activation processes.

The release of cytochrome c into the cytosol of overloaded cardiomyocytes may trigger and accelerate the transition from compensating hypertrophy to right-sided heart failure. Cytosolic cytochrome c can oxidise NADH produced in glycolysis and carry electrons to complex IV of the electron transport chain, bypassing proton pumping by complexes I and III (La Piana et al., 2005; Ripple et al., 2010). The transfer of electrons from cytosolic NADH to cytochrome c may be limited by the cytosolic cytochrome c concentration and will produce heat and reduce ATP/O_2_. Loss of cytochrome c from the intermembrane space impairs the transfer of electrons from complex III to IV (Kuznetsov et al. 2004), and stimulates formation of reactive oxygen species (Zhao et al. 2003). Cytosolic cytochrome c can also induce myocyte apoptosis, which has been demonstrated in end-stage failure (Ecarnot-Laubriet et al. 2002; Campain et al. 2009).

Oxidation of cytosolic NADH by cytochrome c may also inhibit NADH oxidase associated with the ryanodine receptor (RyR2), which regulates Ca^2+^-induced Ca^2+^-release. Low cytosolic NADH/NAD^+^ increases sarcoplasmic reticulum Ca^2+^ release (Zima et al. 2004; Cherednichenko et al. 2004). The diastolic Ca^2+^ concentration is increased and Ca^2+^ transients are delayed during the compensatory phase in MCT-PH (Power et al. 2018). Sabourin et al. (2018) showed that action potential duration, sarcoplasmic reticulum calcium content and Ca^2+^ transients are increased in MCT-PH myocytes from failing hearts.

Myocardial efficiency can also decrease because cross-bridges generate less work/ATP, increased proton permeability of the mitochondrial inner membrane, or increased production of reactive oxygen species. In the latter cases, both activation processes and cross-bridge cycling would require more oxygen than normal. These possibilities are unlikely in view of the heat measurements of Pham et al. (2018), demonstrating normal cross-bridge efficiency in hypertrophied trabeculae.

We tested the hypothesis that mechanical efficiency of isolated papillary muscle is inversely related to the cytosolic cytochrome c concentration. Work and oxygen consumption of papillary muscle preparations dissected from the right ventricle of control and MCT-PH rats were determined using a work loop protocol resulting in maximal power in control papillary muscle subjected to sinusoidal movement (Layland et al. 1995). Blebbistatin was used to discriminate between oxygen consumption for cross-bridge cycling and activation processes allowing confirmation of the results of Pham et al. (2018). Histochemistry of the experimental muscles and the right-sided myocardium was used to determine the cytosolic cytochrome c concentration, the degree of hypertrophy and succinate dehydrogenase (SDH) activity. Proton permeability of the mitochondrial inner membrane, monoamine oxidase (MAO)-A activity and a marker of cardiolipin metabolism were determined in the right-sided myocardium. Glucose 6-phosphate dehydrogenase (G-6-PDH) activity was determined to estimate non-phosphorylating oxygen consumption related to the oxidation of NADPH.

## Methods

Male Wistar rats were obtained from Harlan (n = 6, 2 controls Horst, The Netherlands), or Charles River (n= 17, 6 controls, Sulzfeld, Germany). The local Committee for Animal Experiments (DEC) approved the experiments (Fys 09-14A1V1). The rats were housed in pairs under controlled conditions (temperature 21 to 22°C, humidity 45 to 65% and 12/12 h light dark cycle). They had access to drinking water and food *ad libitum* (Global 2016, Harlan Teklad, Madison, WI). One week after arrival, the experimental rats (body mass 171 to 203 g) were injected subcutaneously with 60 mg monocrotaline in saline/kg body mass (Sigma-Aldrich, St Louis, MO) to induce pulmonary hypertension. Monocrotaline is converted in hepatocytes to monocrotaline pyrrol, which is taken up by erythrocytes at low oxygen tension and released in the pulmonary vasculature during oxygenation, damaging the local endothelium (Xiao et al. 2019). Control rats were not injected. The rats were weighted daily during the week before the efficiency experiments, 22 to 24 days after the injection, when hypertrophy ends and heart failure develops (des Tombe et al. 2002; van Beek-Harmsen et al. 2011). Heart function was assessed by echocardiography under 2-3% isoflurane anaesthesia as described (Hardziyenka et al. 2006; Handoko et al. 2009) on the day before the efficiency experiment.

On the experimental day the rat was anesthetized with isoflurane. The thorax was opened and the beating heart and the lungs were excised and transferred to Tyrode solution cooled on ice (in mM: NaCl 120, KCl 5, CaCl_2_ 1, NaHCO_3_ 27, Na_2_HPO_4_ 2, MgSO_4_ 1.2, glucose 10, 2,3-butanedione monoxime (BDM) 20, N-acetyl cysteine 10, equilibrated with 95% O_2_ and 5% CO_2,_ pH 7.4), in 1.2 ± 0.2 min (mean ± S.D.). N-acetyl cysteine was added as a precaution to scavenge hydroxyl radicals (Aruoma et al. 1989) because trace iron in the Tyrode solution could not be prevented (Reardon & Allen 2009). The heart was transferred to a dissection chamber and perfused via the aorta with Tyrode solution at 10°C in 2.8 ± 0.5 min. Time from opening the chest to perfusion was 4.1 ± 0.7 min. The trachea and large pulmonary vessels were removed and the blotted lung wet and dry mass were determined. After 10 min of perfusion, the right ventricular wall was carefully isolated and frozen in liquid nitrogen. The longest papillary muscle was dissected for the efficiency determination to minimise the unavoidable fraction of damaged myocytes. Muscles were trimmed using fine scissors when they were too thick to be fully oxygenated (Wong et al. 2010) or to optimize the cross-sectional area of the muscle along its length (Kiriazis & Gibbs, 1995). The muscle was transferred to a flow-through chamber at room temperature (22°C) in which the larger and smaller diameters at three locations were measured at 50× magnification using an ocular scale. Muscle volume was calculated assuming elliptical cross-sections. Rings, made from 50 µm diameter platinum wire, were tied to the tendon as close as possible to the myocytes and at the base of the muscle. The apex of the heart and the right ventricular free wall were frozen in liquid nitrogen and stored at -80°C for comparison and additional assays (MAO-A activity: van Eif et al. 2014; proton permeability of mitochondrial inner membrane: Peters et al. 2019).

### Mechanical efficiency of papillary muscle

The papillary muscle was transferred to the oxygen chamber described in detail previously (Wong et al. 2010). The base was connected to the pivot of the stainless steel spinner and the ring attached to the tendon was hooked onto a tungsten wire, which was suspended from a force transducer (SensoNor AE-801, Horten, Norway) via a capillary. The force transducer was connected to a servomotor. Tyrode solution without BDM containing 1.5 mM Ca^2+^, equilibrated with 95%O_2_ and 5%CO_2_ at 37°C, was pumped into the chamber. The oxygen tension (PO_2_) in the chamber was determined using a polarographic oxygen electrode. After stabilization the pump was stopped, and PO_2_ in the chamber decreased steadily. The decrease was due to basal oxygen consumption, oxygen consumption by the electrode and oxygen loss to the surroundings. The rate at which PO_2_ decreased without a preparation in the chamber varied between experiments, hampering an accurate estimate of the basal rate of oxygen consumption of the preparation. Mechanical efficiency of the papillary muscle was calculated assuming all suprabasal oxygen uptake is used to oxidise glucose (Woledge et al. 1985, p.209): efficiency = work/(suprabasal oxygen consumption x 473kJ/mol O_2_), where 473 kJ/mol O_2_ is the heat of combustion of glucose (Carpenter, 1939).

### Protocol

After equilibration at 37°C, the stimulus threshold was determined and a force-length relationship was determined at 0.5 Hz, 0.2 ms stimuli 20% above threshold, by increasing the length of the preparation in steps of 100 or 50 µm. Contraction time, relaxation time and tension were determined. After determination of the length giving maximum force (L_max_), the length of the muscle was reduced to L_opt =_ 0.925 L_max_. Sinusoidal length changes were imposed on the muscle with peak-to-peak amplitude 0.15 L_max_ (Layland et al. 1995). The frequency was 5 Hz, similar to the heart rate during sleep (Kramer et al. 1995). The stimulus phase giving maximum work was used. The conduction delay of the stimulus was determined during the efficiency run from the time between the upstroke of the stimulus pulse to the start of force development.

A typical efficiency determination lasted for 10 min: 3 min without stimulation, 4 min 5 Hz and 3 min recovery. After each run the Tyrode’s solution in the chamber was replaced at a rate of 12 ml/min for 20 min.

To investigate whether muscles became hypoxic during the efficiency run, isoprenaline was injected into the experimental chamber after the efficiency experiment (1 µM final concentration; 3 control and 2 MCT-PH muscles).

After washout, blebbistatin (Calbiochem 203389, 50mM in 90% dimethyl sulfoxide; Merck, Amsterdam) was injected into the chamber at 10 µM final concentration. After 10 min irreversible inhibition of active force was achieved and Tyrode’s solution without blebbistatin was pumped into the chamber. The efficiency protocol was repeated to determine oxygen consumption for activation processes (Pham et al. 2017).

After determination of the effect of blebbistatin the protocol was repeated in 6 MCT-PH muscles in Tyrode solution containing 50 µM ruthenium red to determine the effect of calcium cycling on oxygen consumption (Zhou & Bers, 2002). A final run was started after 30 min of washout of ruthenium red to determine recovery.

After opening the oxygen chamber, L_opt_ was determined and the muscle was embedded in Tyrode’s solution containing 15% gelatine (pH 7.4). After solidification on ice, the preparation was frozen in liquid nitrogen and stored at -80°C. In 14 muscles the volume the muscle was determined again under the dissection microscope before embedding. The volume determination was accurate: muscle volume after experiment = 0.84 (standard error 0.03) × volume before (F_1,12_ = 990.2, P < 0.0001, regression line forced through the origin).

### Data analyses

The oxygen electrode current, motor position, force and stimulus pulse were digitized and sampled at 1 kHz by Amsterdam UMC DAQ, a custom-written, LabVIEW (National Instruments Netherlands BV, Woerden, NL) based data acquisition program. Force as a function of length was displayed and work per cycle was calculated real-time from the sum of the products of force and length change.

The decay of the oxygen tension in the chamber before and after the stimulation period was used to construct a second order polynomial, which was subtracted from the recording to determine suprabasal oxygen consumption due to stimulation of the preparation (Elzinga & van der Laarse, 1988). The volume of the oxygen chamber was 381 µl. The solubility of oxygen at 760 mmHg and 37°C in Tyrode’s solution used was 22.73 μl O_2_/ml (Bartels et al. 1971). Oxygen consumption was normalized by the volume of the preparation determined before mounting. Microsoft Excel and Amsterdam UMC DAQ Analyzer were used for analyses.

### Histochemistry, morphometry and microdensitometry

Cryostat sections, 5 µm thick, were cut at -20°C from the experimental muscle and for comparison from the right-sided free wall or the apex of the heart. Because the number of sections that could be cut from the experimental muscle was limited, we incubated for succinate dehydrogenase (SDH, complex II) activity and cytosolic cytochrome c. The activity of SDH determined at 660 nm is proportional to the maximum rate of oxygen consumption in single *Xenopus* muscle fibres (van der Laarse et al. 1989) and control myocardial trabeculae (des Tombe et al. 2002). The sections were also used to measure interstitial space and damaged myocytes (van der Laarse et al. 2005) – cells which had lost SDH activity.

Cytosolic cytochrome c was determined using calibrated immunohistochemistry in the experimental muscle and the right-sided free wall as described (van Beek-Harmsen & van der Laarse, 2005). The absorbance of the final diaminobenzidine precipitate was determined at 436 nm.

The histochemical assays in the experimental muscles were extended with activities of complex V in the right ventricular free wall, which includes the determination of proton permeability of the mitochondrial inner membrane (IMM PP). A modification of Meijer & Vloedman’s (1980) method for mitochondrial coupling was used to determine F_1_F_o_ATPase (complex V) activity in the right ventricular free wall. Maximum activity is obtained in the presence of 1 mM 2,4-dinitrophenol (DNP) which carries protons across the inner mitochondrial membrane. Background activity is measured after inhibition of complex V by 25 µM oligomycin. The absorbance of the final PbS precipitate was determined at 550 nm. The modified method is quantitative (Peters et al. 2019).

Glucose 6-phosphate dehydrogenase (G-6-PDH) activity was determined at 37^°^C in 7 µm thick sections of the right ventricular free wall (van Noorden, 1984; van Noorden & Frederiks, 1992). This method allows an estimate of maximum rate of NADPH production in the pentose phosphate pathway. The activity of G-6-PDH was calculated from the increase in absorbance between 15 min and 30 min incubation time, corrected for background activity measured without NADP, using an extinction coefficient of 19000 M^-1^cm^-1^ at 550 nm.

MAO activity was demonstrated using a modification of the quantitative method described by Frederiks & Marx (1985). The sections were incubated in oxygenated 25 mM sodium phosphate buffer, pH 7.7, 6.25 mM tryptamine HCl, and 0.4 mM tetranitroblue tetrazolium. The absorbance of the formazan precipitate in 5 µm thick sections is proportional to incubation time and was determined after 90 min incubation at 37^°^ at 550 nm. Sections preincubated for 20 min with 2 µM of the irreversible MAO-A inhibitor clorgyline did not stain.

The absorbance of the final reaction products was determined using a calibrated microdensitometer (Lee-de Groot et al. 1998). To obtain a reliable estimate of the mean absorbance and myocyte cross-sectional area, six to thirty myocytes, cut perpendicularly to the longitudinal axis, were measured. Images of the sections were analysed using NIH Image or ImageJ (http://rbs.info.nih.gov), taking the pixel aspect ratio into account.

### Phosphatidylglycerol and cardiolipin

Phosphatidylglycerol and cardiolipins were analyzed in the right-sided myocardium by high performance liquid chromatography-mass spectrometry (Houtkooper et al. 2009). The ratio of the most abundant phosphatidylglycerol (PG34:1, a precursor of cardiolipin) and tetralinoleylcardiolipin (CL 72:8) was used as indicator of cardiolipin metabolism. The data on PG 34:1/CL 72:8 were related to proton permeability of the mitochondrial inner membrane in a previous report (Peters et al. 2019).

### Subdivision of oxygen uptake

An estimate of basal oxygen consumption was unreliable in the present experiments because determinations of oxygen loss from the chamber gave variable results. Therefore, only suprabasal oxygen consumption is reported.

Oxygen uptake for activation processes and for cross-bridge cycling are:

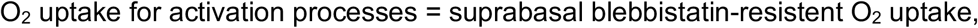

and:

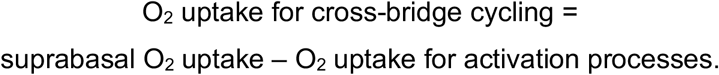

Oxygen consumption for mitochondrial calcium cycling was estimated by additional inhibition experiments using ruthenium red to inhibit mitochondrial calcium influx after blebbistatin inhibition.

Proton permeability of the inner mitochondrial membrane (IMM PP) relative to the permeability with DNP was determined:

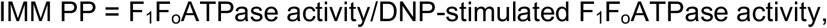

where the ATPase activities are determined by quantitative enzyme histochemistry as described above. Increased IMM PP will increase oxygen consumption to compensate for proton leak and reduce ATP/O_2_, but the relationship is not proportional (for discussion, see Scheffler, 2003; Korzeniewski, 2017).

G-6-PDH activity in sections of the RV wall was determined to estimate oxygen consumption related to the oxidation of NADPH. NADPH produced in the pentose phosphate pathway is an important H_2_O_2_ scavenger. NADPH reduces glutathione disulphide (GSSG → 2 GSH, corresponding to 1 O_2_/2 NADPH), and is a substrate for nitric oxide production by nitric oxide synthase (1.5 O_2_/NADPH) and superoxide production by NADPH oxidase (2 O_2_/NADPH). In addition, the pentose phosphate pathway produces ribulose 5-phosphate, which is required for increased xanthine oxidase activity or re-enters glycolysis via the non-oxidative phase of the pentose phosphate pathway. Two NADPH molecules and one ribulose 5-phosphate are produced per glucose 6-phosphate (Salway, 2017). We assume that at most 4 O_2_ is required for non-mitochondrial oxidation of the products of 1 glucose 6-phosphate in the pentose phosphate pathway.

### Statistics

Values are mean ± SD unless indicated otherwise. Student’s two-sided t-test, with equal or unequal variance where appropriate, was used to determine differences between groups (Microsoft Excel). Prism 9.1.0 (Graphpad, San Diego, CA) was used for further statistical analyses. Unless indicated otherwise, relationships between two measured variables were determined using Pearson correlations and Deming regression analyses on homogenised and dimensionless data, obtained after transformation: x_i_’ = (x_i_ – Σx_i_/n)/SD, making mean x’ = 0, and variance = 1. The exact probability or P < 0.0001 is reported, P < 0.05 is taken as significant.

## Results

Table 1 shows characteristics of the rats. Body mass of control rats increased whereas body mass of the MCT-PH rats decreased during the days before the efficiency measurement. The body mass change correlated non-linearly with cardiac output (Spearman rank correlation coefficient r_s_ = 0.72, P=0.0002), stroke volume (r_s_ = 0.75, p=0.0001), and pulmonary artery flow acceleration time (PAAT, r_s_ = 0.72, P=0.0002; PAAT/cycle length, r_s_ = 0.66, P = 0.0011, which are inversely related to pulmonary artery systolic pressure; Jones et al. 2002; Handoko et al. 2009).

**Table 1.** Characteristics of the rats and RV free wall.

| Group | Control |  | MCT-PH |  | P |
| --- | --- | --- | --- | --- | --- |
| Item | mean ± SD (range) | n | mean ± SD (range) | n | |
| Body mass (g) | 300 ± 16 (276 - 308) | 8 | 259 ± 13 (241 - 287) | 15 | <0.0001 |
| Body mass change (%/day) | 1.2 ± 1.1 (-0.3 - 2.6) | 6 | -0.9 ± 1.3 (-4.0 - 0.8) | 14 | 0.0039 |
| Lungs wet mass (g) | 1.34 ± 0.12 (1.23 - 1.54) | 8 | 2.13 ± 0.29 (1.50 - 2.64) | 15 | <0.0001 |
| Cardiac output (ml/min) | 96 ± 15 (78 - 116) | 8 | 56 ± 19 (34 - 97) | 15 | <0.0001 |
| Stroke volume (ml) | 0.24 ± 0.05 (0.20 - 0.34) | 8 | 0.16 ± 0.05 (0.09 - 0.24) | 15 | 0.0012 |
| Heart rate (beats/min) | 400 ± 30 (345 - 432) | 8 | 354 ± 27 (408 - 326) | 15 | 0.0028 |
| PAAT (ms) | 25 ± 6 (19 - 36) | 8 | 15 ± 2 (12 - 18) | 15 | 0.0016 |
| PAAT/cycle length (%) | 17 ± 4 (15 - 25) | 8 | 9 ± 1 (7 - 11) | 15 | 0.0004 |
| TAPSE (mm) | 2.9 ± 0.6 (2.1 - 3.9) | 8 | 2.0 ± 0.6 (1.3 - 3.2) | 15 | 0.0040 |
| RV wall thickness (mm) | 0.95 ± 0.01 (0.8 - 1.1) | 8 | 1.31 ± 0.15 (0.9 - 1.4) | 15 | <0.0001 |
| RV PG34:1/CL72:8 | 0.017 ± 0.004 (0.014 - 0.022) | 4 | 0.035 ± 0.011 (0.019 - 0.050) | 11 | 0.0059 |
| RV G-6-PDH (μM/s) | -0.1 ± 0.5 (-0.2 - 0.6) | 7* | 2.8 ± 1.3 (0.6 - 4.6) | 13 | <0.0001 |
| RV MAO-A ($A_{550nm}$) | 0.21 ± 0.06 (0.15 - 0.28) | 8 | 0.30 ± 0.10 (0.15 - 0.49) | 15 | 0.0075 |
| RV SDH ($A_{660nm}$) | 0.13 ± 0.04 (0.09 - 0.19) | 7 | 0.13 ± 0.04 (0.05 - 0.21) | 14 | 0.9390 |
| RV $F_1F_0$ATPase + DNP ($A_{550nm}$) | 0.27 ± 0.16 (0.09 - 0.52) | 8 | 0.23 ± 0.09 (0.08 - 0.35) | 15 | 0.4965 |
| RV cytosolic cyt c (μM) | 14 ± 12 (-3 - 27) | 5*<sup>†</sup> | 37 ± 39 (31 - 129) | 15 | 0.0693 |

\*excluding one outlier (open circle in Figs. 1,3 and 4, see text). <sup>†</sup>sections were unavailable for two experiments. Most (17/23) MAO-A activities have been reported by van Eif et al. (2014).

Weak correlations were found between body mass change and right ventricular wall thickness (r_s_ = -0.47, P = 0.0290), tricuspid annular plane systolic excursion (TAPSE, r_s_ = 0.50, P = 0.0184) and heart rate (r_s_ = 0.42, P = 0.0523). The change of body mass correlated inversely with wet mass of the lungs (r_s_ = -0.70, P = 0.0003). Pleural effusion was not observed. Dry and wet mass of the lungs were proportional: dry mass = 0.190 (standard error 0.002) x wet mass (F_1,14_ = 7086, p <0.0001, regression line forced through the origin). The increase in PG/CL was due to both an increase of PG34:1 from 0.87 ± 0.09 nmol/mg protein (n = 3) to 1.19 ± 0.14 nmol/mg (n = 7; P = 0.0080) and a decrease of CL72:8 from 52 ± 9 nmol/mg to 31 ± 8 nmol/mg (P = 0.0072).

The activities of RV SDH and RV F_1_F_o_ATPase (+DNP) ranged widely within groups, but were similar in MCT-PH and control. These complex activities correlated weakly (r = 0.56, P = 0.0076). Cytosolic cytochrome c in RV cardiomyocytes also ranged widely, from undetectable in both groups to 163 µM in the outlier indicated by the open circle in Fig. 2*DHLP*.

The activity of G-6-PDH was undetectable in 6/7 (Fig. 2*BFJN*) control hearts, but was increased in the control rat which stopped breathing during anesthesia. The heart was beating when it was excised (see Discussion).

G-6-PDH activity was increased in the RV free walls of all MCT-PH rats (to 2.8 µM glucose 6-phosphate/s, Table 1), and was an order of magnitude higher in clusters of inflammatory cells, mainly located at the junction of the RV wall and the septum (Fig. 1). Assuming 1 to 4 O_2_ are required to oxidise the products of glucose 6-P turnover in the pentose phosphate pathway (see Methods), the rate of oxygen consumption due to the production and/or scavenging of reactive oxygen in MCT-PH cardiomyocytes is 2.8 to 11.2 µM/s, respectively, or 5 to 19% of suprabasal oxygen consumption during the efficiency runs in MCT-PH rats (Table 2).

**Figure 1.**
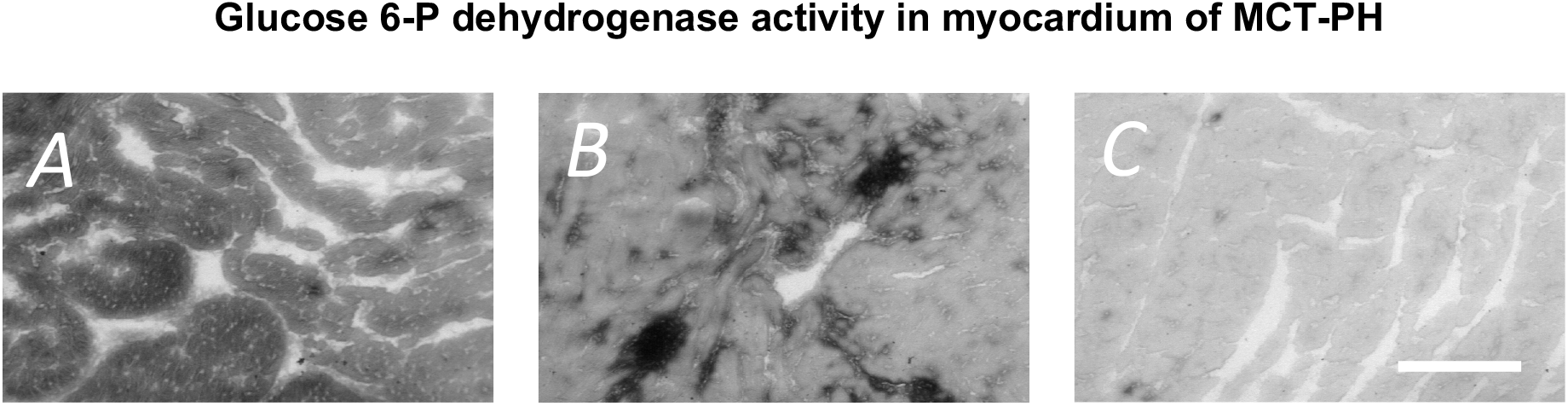
A: right-sided myocardium, B: septum, high activity (dark) clusters are inflammatory cells, C: left-sided myocardium, where activity is similar to background activity in RV and septum. Scale bar 100 µm.

**Table 2.**
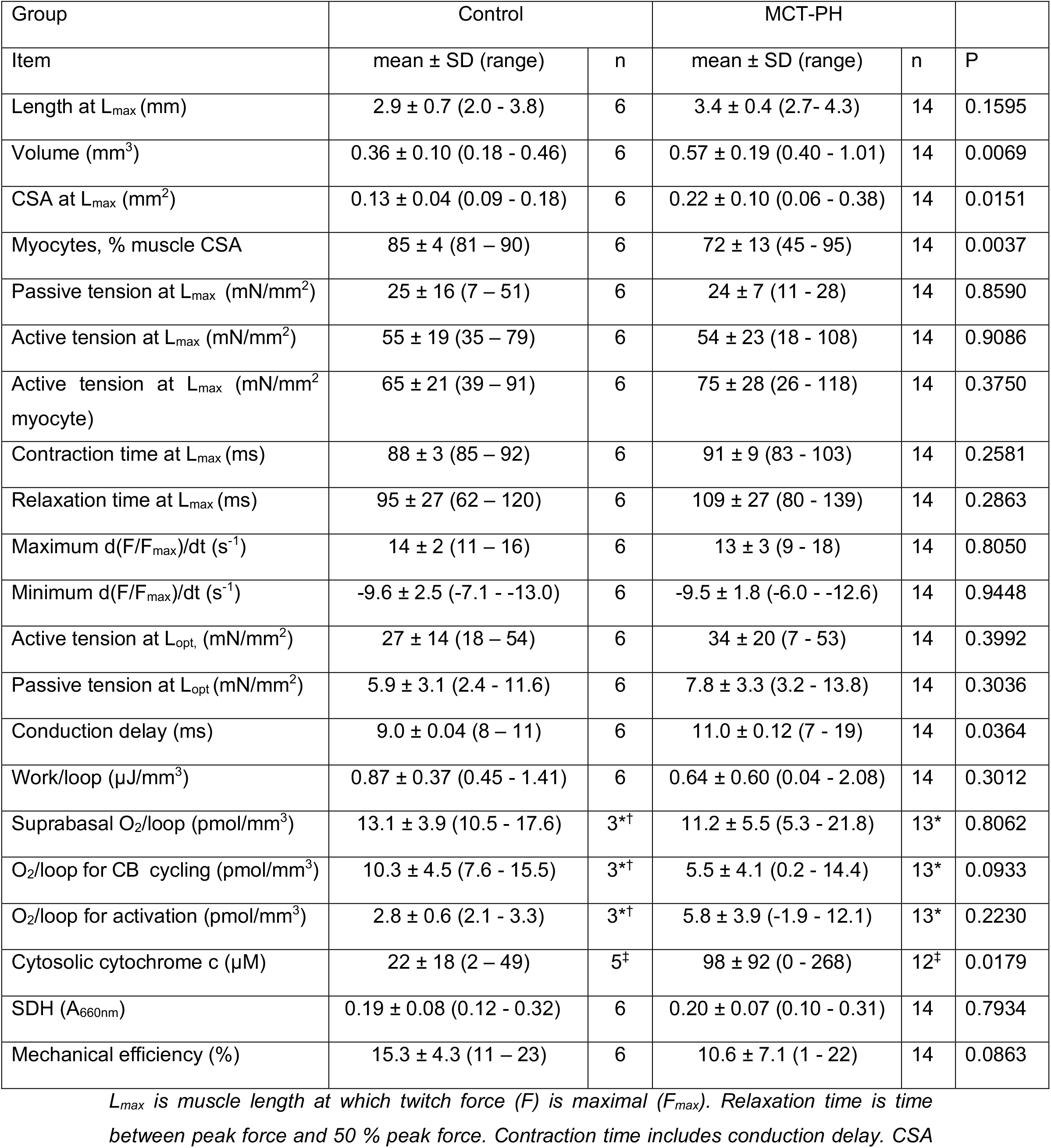
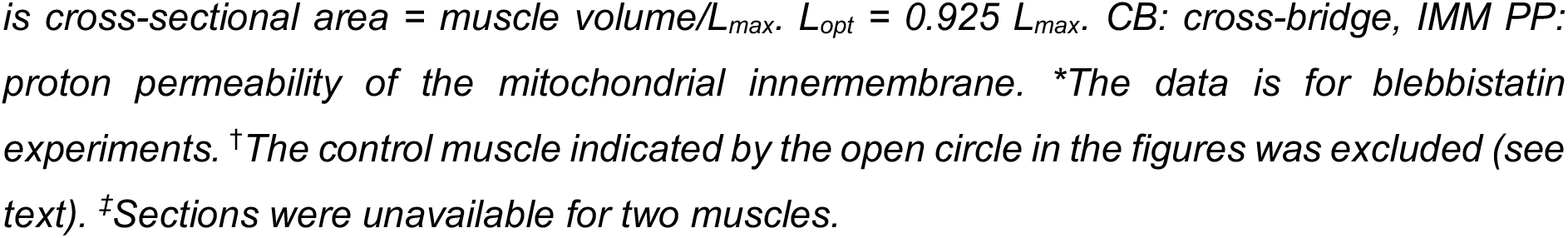
Characteristics of experimental papillary muscles.

| Group | Control |  | MCT-PH |  |  |
| --- | --- | --- | --- | --- | --- |
| Item | mean ± SD (range) | n | mean ± SD (range) | n | P |
| Length at $L_{\max}$ (mm) | 2.9 ± 0.7 (2.0 - 3.8) | 6 | 3.4 ± 0.4 (2.7 - 4.3) | 14 | 0.1595 |
| Volume (mm<sup>3</sup>) | 0.36 ± 0.10 (0.18 - 0.46) | 6 | 0.57 ± 0.19 (0.40 - 1.01) | 14 | 0.0069 |
| CSA at $L_{\max}$ (mm<sup>2</sup>) | 0.13 ± 0.04 (0.09 - 0.18) | 6 | 0.22 ± 0.10 (0.06 - 0.38) | 14 | 0.0151 |
| Myocytes, % muscle CSA | 85 ± 4 (81 – 90) | 6 | 72 ± 13 (45 - 95) | 14 | 0.0037 |
| Passive tension at $L_{\max}$ (mN/mm<sup>2</sup>) | 25 ± 16 (7 – 51) | 6 | 24 ± 7 (11 - 28) | 14 | 0.8590 |
| Active tension at $L_{\max}$ (mN/mm<sup>2</sup>) | 55 ± 19 (35 – 79) | 6 | 54 ± 23 (18 - 108) | 14 | 0.9086 |
| Active tension at $L_{\max}$ (mN/mm<sup>2</sup> myocyte) | 65 ± 21 (39 – 91) | 6 | 75 ± 28 (26 - 118) | 14 | 0.3750 |
| Contraction time at $L_{\max}$ (ms) | 88 ± 3 (85 – 92) | 6 | 91 ± 9 (83 - 103) | 14 | 0.2581 |
| Relaxation time at $L_{\max}$ (ms) | 95 ± 27 (62 – 120) | 6 | 109 ± 27 (80 - 139) | 14 | 0.2863 |
| Maximum d(F/$F_{\max}$)/dt (s<sup>-1</sup>) | 14 ± 2 (11 – 16) | 6 | 13 ± 3 (9 - 18) | 14 | 0.8050 |
| Minimum d(F/$F_{\max}$)/dt (s<sup>-1</sup>) | -9.6 ± 2.5 (-7.1 - -13.0) | 6 | -9.5 ± 1.8 (-6.0 - -12.6) | 14 | 0.9448 |
| Active tension at $L_{\text{opt}}$ (mN/mm<sup>2</sup>) | 27 ± 14 (18 – 54) | 6 | 34 ± 20 (7 - 53) | 14 | 0.3992 |
| Passive tension at $L_{\text{opt}}$ (mN/mm<sup>2</sup>) | 5.9 ± 3.1 (2.4 - 11.6) | 6 | 7.8 ± 3.3 (3.2 - 13.8) | 14 | 0.3036 |
| Conduction delay (ms) | 9.0 ± 0.04 (8 – 11) | 6 | 11.0 ± 0.12 (7 - 19) | 14 | 0.0364 |
| Work/loop (μJ/mm<sup>3</sup>) | 0.87 ± 0.37 (0.45 - 1.41) | 6 | 0.64 ± 0.60 (0.04 - 2.08) | 14 | 0.3012 |
| Suprabasal O<sub>2</sub>/loop (pmol/mm<sup>3</sup>) | 13.1 ± 3.9 (10.5 - 17.6) | 3<sup>*†</sup> | 11.2 ± 5.5 (5.3 - 21.8) | 13<sup>*</sup> | 0.8062 |
| O<sub>2</sub>/loop for CB cycling (pmol/mm<sup>3</sup>) | 10.3 ± 4.5 (7.6 - 15.5) | 3<sup>*†</sup> | 5.5 ± 4.1 (0.2 - 14.4) | 13<sup>*</sup> | 0.0933 |
| O<sub>2</sub>/loop for activation (pmol/mm<sup>3</sup>) | 2.8 ± 0.6 (2.1 - 3.3) | 3<sup>*†</sup> | 5.8 ± 3.9 (-1.9 - 12.1) | 13<sup>*</sup> | 0.2230 |
| Cytosolic cytochrome c (μM) | 22 ± 18 (2 – 49) | 5<sup>‡</sup> | 98 ± 92 (0 - 268) | 12<sup>‡</sup> | 0.0179 |
| SDH (A<sub>660nm</sub>) | 0.19 ± 0.08 (0.12 - 0.32) | 6 | 0.20 ± 0.07 (0.10 - 0.31) | 14 | 0.7934 |
| Mechanical efficiency (%) | 15.3 ± 4.3 (11 – 23) | 6 | 10.6 ± 7.1 (1 - 22) | 14 | 0.0863 |

$L_{\max}$ is muscle length at which twitch force (F) is maximal ( $F_{\max}$ ). Relaxation time is time between peak force and 50 % peak force. Contraction time includes conduction delay. CSA
*is cross-sectional area = muscle volume/ $L_{max}$ . $L_{opt} = 0.925 L_{max}$ . CB: cross-bridge, IMM PP: proton permeability of the mitochondrial innermembrane. \*The data is for blebbistatin experiments. †The control muscle indicated by the open circle in the figures was excluded (see text). ‡Sections were unavailable for two muscles.*

### Glucose 6-P dehydrogenase activity in myocardium of MCT-PH

Several characteristics of MCT-PH rats in Table 1 overlapped with controls, indicating that the effect of MCT was rather variable, however, most differences between the groups were highly significant, and confirm right-sided heart failure in MCT-PH rats.

Fig. 2 shows the relationships between four of the echocardiography characteristics related to hypertrophy and work and the marker of cardiolipin metabolism (phosphatidylglycerol/cardiolipin, PG/CL), glucose-6-phosphate dehydrogenase activity, MAO-A activity and cytosolic cytochrome c, all determined in the right ventricular free wall. Significant Deming regressions were found, except for heart rate and PG/CL (Fig. 2*I*), and the relationships with cytosolic cytochrome c (Fig. 2*DHLP*). Despite considerable variation, subgroup regressions were only significant for wall thickness and G-6-PDH (Fig 1*B*, P = 0.0319) and PAAT/CL and MAO-A (Fig 1*O*, P = 0.0122) in MCT-PH.

**Figure 2.**
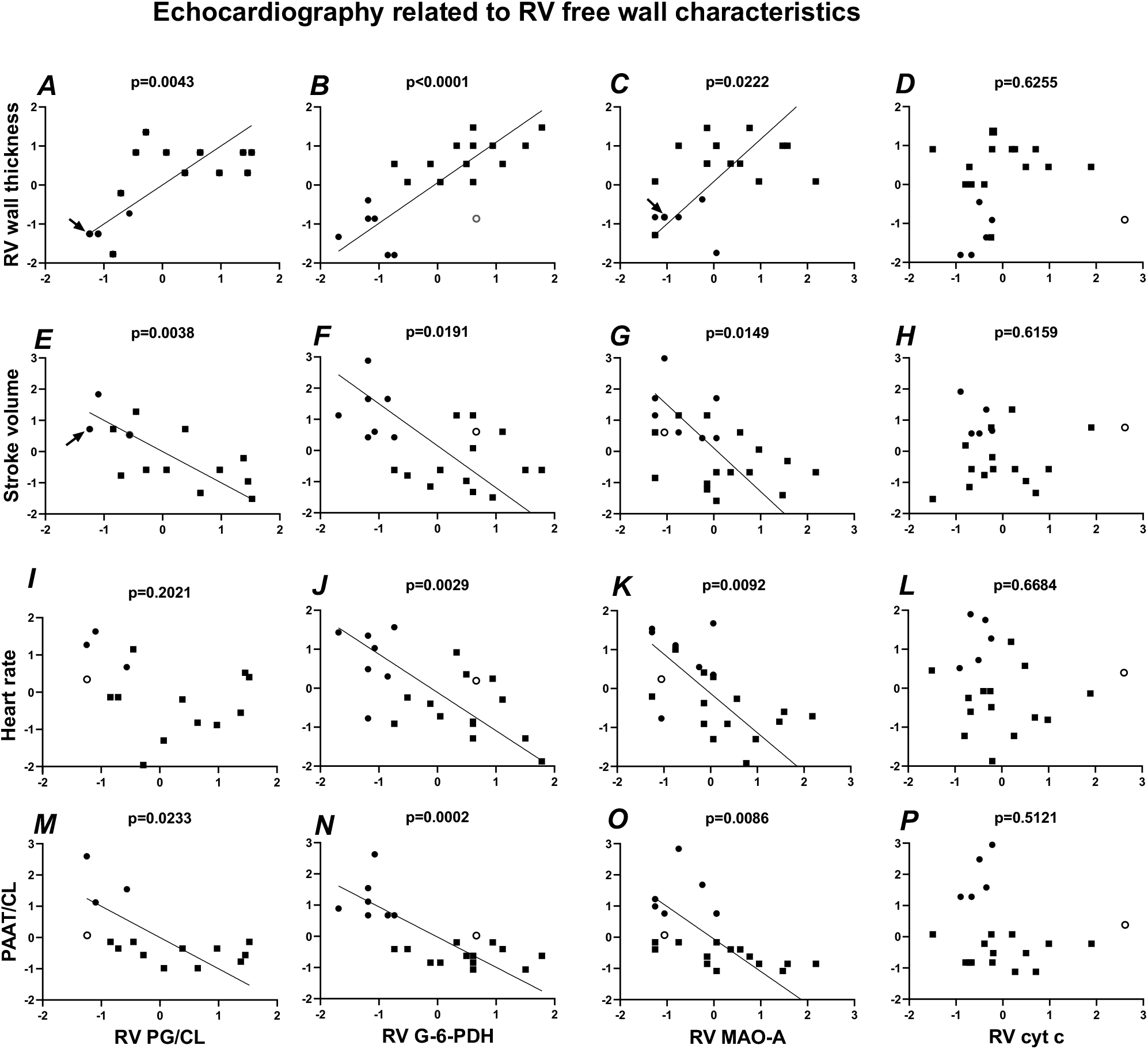
Deming regression analyses of homogenised and dimensionless values of echocardiography and RV myocardial characteristics. Circles: control, Squares: MCT-PH. The arrows point at overlapping values, including the control RV myocardium indicated by the open circle (see text) in C. P values indicate significance of Deming regressions. PG/CL is phosphatidylglycerol 34:1/cardiolipin 72:8. Table 1 gives untransformed data per experimental group.

### Characteristics of papillary muscles

After mounting the preparation in the oxygen chamber, the force-length relationship was determined at 0.5 stimuli/s, 37°C and 1.5 mM Ca^++^. Twitch characteristics were determined at the length giving maximum force (L_max_, Table 2). Passive and active tensions of control and MCT-PH at L_max_ were similar. The conduction delay was 2 ms longer in MCT-PH during efficiency determinations, but other twitch characteristics were not significantly different.

### Efficiency determination

In 20/23 hearts (6 control, 14 MCT) efficiency measurements were successful. Two control and one MCT preparation contracted out of phase during a part of the stimulation period. These muscles are excluded from efficiency analyses.

Fig. 3*A-C* shows work loops of a control muscle and two extreme examples of MCT-PH muscles. Passive loops - without stimulation - are relatively small and proceed clockwise. The area of these loops corresponds to viscoelastic energy loss associated with passive stretch-relaxation cycles. Loops during which positive work is produced are larger and proceed counter-clockwise. Maximum work per loop is produced when the muscles are stimulated during stretch (Layland et al, 1995): at 34 ± 6 degrees in control (31 ± 9 ms before maximum length is reached) and 27 ± 9 degrees (35 ± 12 ms before maximum length) in MCT-P (P = 0.1143).

**Figure 3 A-C.**
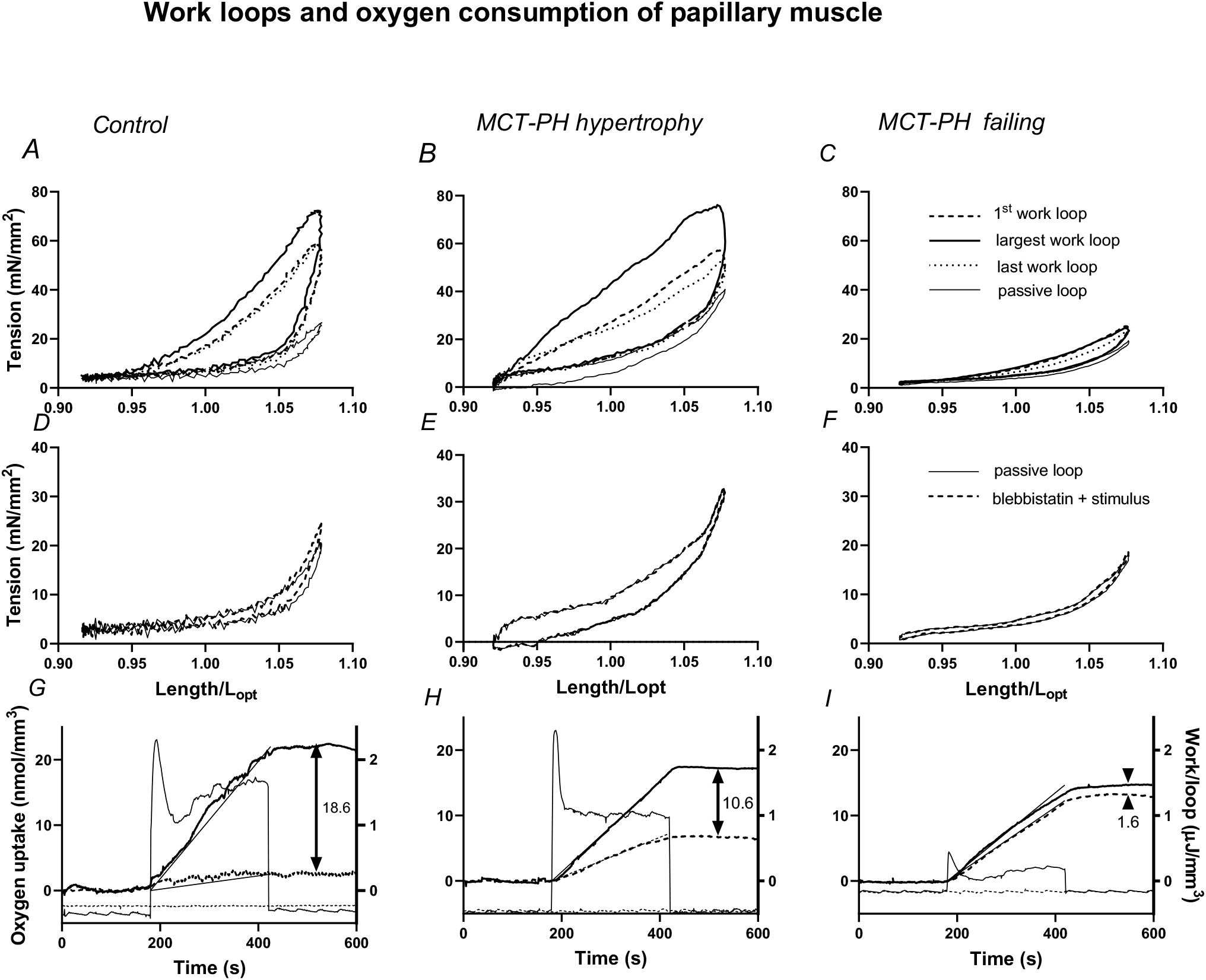
Examples of work loops during sinusoidal length changes for 4 min of 5 Hz stimulation in a control papillary muscle preparation (A) and two extreme examples of MCT-PH papillary muscles (BC). Passive loops (thin curves) are without stimulation - time proceeds clockwise and the area of the loop represents work done by the motor on the muscle. The first, last and largest active work loops with stimulation are also shown. In these loops time proceeds anti-clockwise. The area of the loop is proportional to work done by the muscle. D-F show a passive loop and a loop after blebbistatin inhibition with stimulus. Stimulus phases were: left column 34°, middle 40°, right 26°. G-I show work (thin tracings) and suprabasal oxygen uptake (thick traces) as a function of time. The muscles are stimulated from 180 to 420 s at 5 Hz. Dashed tracings are work and oxygen consumption by stimulated muscles after blebbistatin inhibition. The difference between oxygen uptake before and after blebbistatin inhibition is indicated by arrows and estimates oxygen consumption for cross-bridge cycling. Work and oxygen consumption are normalised by muscle volume and muscle length by L_opt_. The lines through the oxygen uptake tracings connect (180 s, 0) and (420 s, mean total oxygen uptake), for comparison.

Fig. 3*D-F* compares work loops with stimulus after treatment with blebbistatin to passive work loops. These loops are similar, confirming that blebbistatin inhibition in the oxygen chamber is irreversible. Fig. 3*G-H* shows work/loop and suprabasal oxygen uptake used to calculate efficiency. Stimulation at 5 Hz starts at 180 s and stops at 420 s. Work per loop transiently increases and becomes fairly steady during the last minute of stimulation. Comparison with the lines from 180 to 420 s indicates that oxygen uptake increases rapidly and is fairly constant during at least the first 180 s of stimulation. Work done by MCT-PH muscles varied considerably. Mechanical efficiency is calculated from total positive work – the area of the active loops in Fig. 3*A-C* - and suprabasal oxygen consumption.

Work is negative after blebbistatin inhibition, but stimulation induces activation-related oxygen consumption, which increases in MCT-PH muscles, both in absolute and relative amount. In the extreme example shown in Fig 3*I*, activation-related oxygen consumption is 89% of total oxygen consumption before cross-bridge inhibition.

In Fig. 4*A* work of 1200 loops, determined as shown in Fig. 3*G-I*, is plotted against suprabasal oxygen uptake. Mean values are listed in Table 2. Considerable variation was observed for work and oxygen consumption of different muscles and also for efficiency within groups. These muscle characteristics were not related to the cross-sectional area of the muscle or to interstitial space (results not shown). The relationships between work and oxygen consumption were not significantly different for control and MCT-PH muscles (Table 2 and Fig. 4). However, the relationships between efficiency and cardiomyocyte cross-sectional area of control and MCT-PH (Fig. 4*B*) were significantly different. Fig. 4*B* suggests that an optimal myocyte cross-sectional area in rat exists: the regression lines for control and MCT-PH intersect at normalised CSA = -0.346, corresponding to 420 µm^2^. Efficiency in MCT-PH decreased with increasing myocyte cross-sectional area (F_1,11_ = 6.190, P = 0.0301).

**Figure 4.**
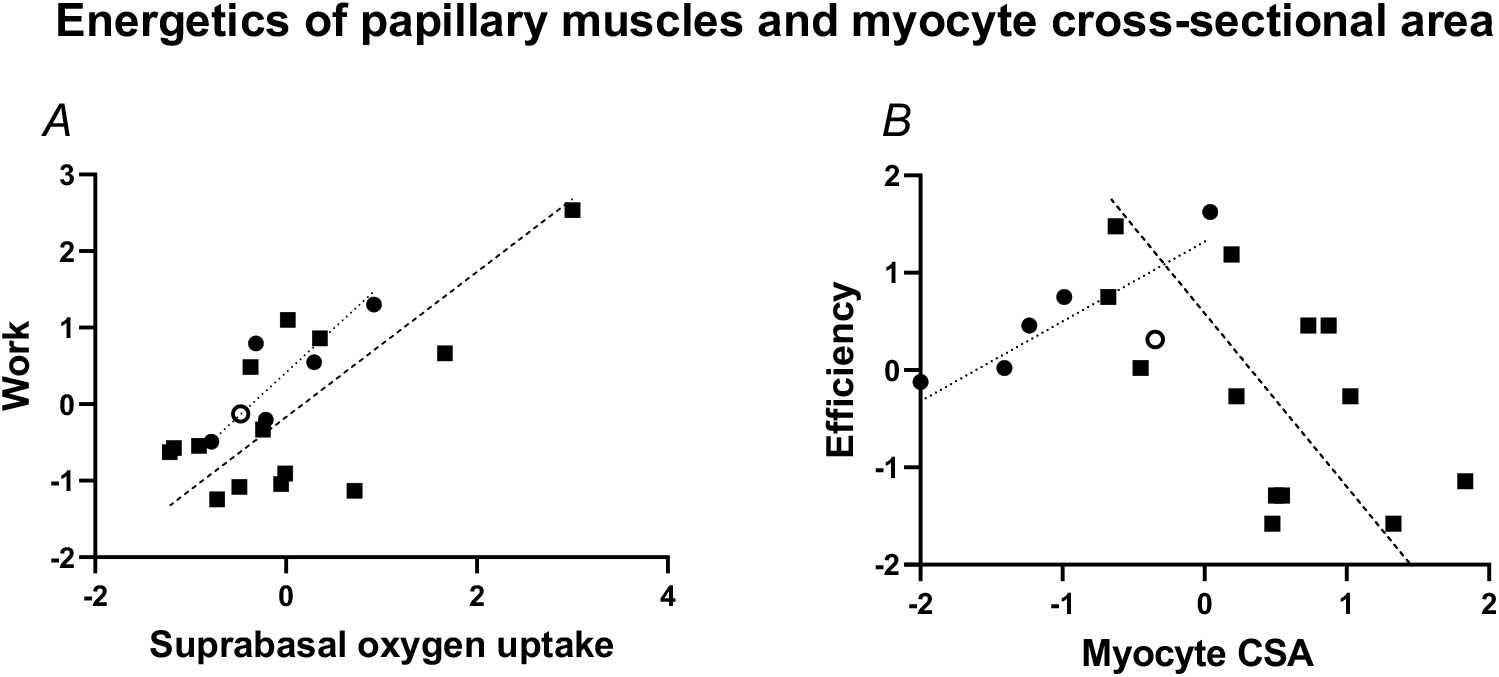
Normalised and homogenised work and oxygen uptake for control and MCT-PH papillary muscles. Deming regression lines are shown; circles and dotted lines: control; squares and interrupted lines: MCT-PH. A: Work plotted against oxygen uptake, the open circle indicates the control muscle dissected from the heart with high G-6-PDH activity (Fig. 2). The regression lines in A are not significantly different (slopes: F_1,16_ = 0.2245, P = 0.6420; intercepts: F_1,17_ = 2.301, P = 0.1477). B: Efficiency plotted against myocyte cross-sectional area (at L_opt_) in cryostat sections of the preparation after the experiment. Efficiency of the muscle is calculated from the data in A. The slopes for control and MCT-PH in B are significantly different (F_1,15_ = 7.390, P = 0.0159).

### Adrenergic stimulation

In three control and two MCT-PH muscles efficiency determinations were repeated in the presence of 1 µM isoprenaline, to see whether oxygen diffusion was a limiting factor in the efficiency experiments. In two control and two MCT-PH muscles, suprabasal oxygen consumption during stimulation doubled in the presence of isoprenaline, excluding oxygen limitation in these preparations during the first efficiency run. Work increased in one of the control muscles (by 86 %), but not in the other (−1%) and decreased in the MCT-PH muscles (−3% and -26%). In the third control muscle oxygen consumption decreased by 27% and work increased by 23%. It cannot be excluded that isoprenaline induced hypoxia in the preparations.

### Determinants of efficiency

Fig. 5 shows images used to determine cytosolic cytochrome c in the experimental muscles. Gelatin sections containing known amounts of cytochrome c were used for calibration. The cytosolic cytochrome c in the muscle myocytes was higher than the cytosolic cytochrome c observed in the RV free wall (compare Tables 1 and 2). There was a strong correlation between cytosolic cytochrome c concentrations in the muscle and the free wall, with two outliers which were muscles exposed to ruthenium red. After excluding these two outliers, cytosolic cytochrome c in the RV myocytes = 0.61 (standard error 0.04) x cytosolic cytochrome c in papillary muscle myocytes (F_1,14_ = 233.3, P < 0.0001; regression line forced through the origin). The cytosolic cytochrome c concentration did not correlate with efficiency, but it correlated with work (Table 3). Work was the most important determinant of efficiency (r^2^ = 0.68). Suprabasal oxygen consumption did not correlate with efficiency. Oxygen consumption after blebbistatin inhibition did not correlate with work or oxygen consumption for cross-bridges but was an independent determinant of efficiency (r^2^ = 0.32), in agreement with the activation heat results of Pham et al. (2018). RV IMM PP was a determinant of efficiency (r^2^ = 0.35). The variance of efficiency is explained by the three significant variables in Table 3 (coefficient of multiple determination R^2^ = 0.79, F_3,13_ = 16.52, P = 0.0001).

**Figure 5.**
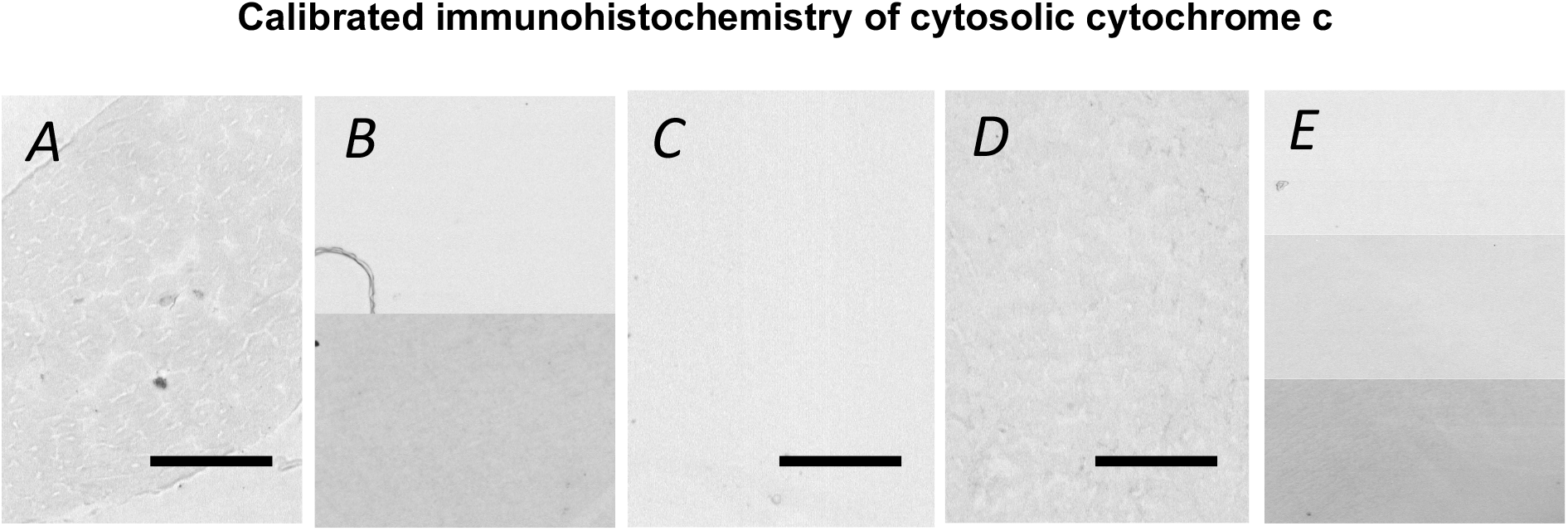
Microdensitometry images at 436 nm. A: control muscle, efficiency 23%; scale bar 50 µm. B: Absorbance calibration for A; top background (no added cytochrome c), bottom 0.15 mM cytochrome c. C: MCT-PH papillary muscle, efficiency = 17 %, D: MCT-PH muscle efficiency = 1%. Scale bars C, D: 100 µm. E: Absorbance calibration for C, D: top background, middle 0.15 mM, bottom 0.3 mM cytochrome c.

**Table 3.**
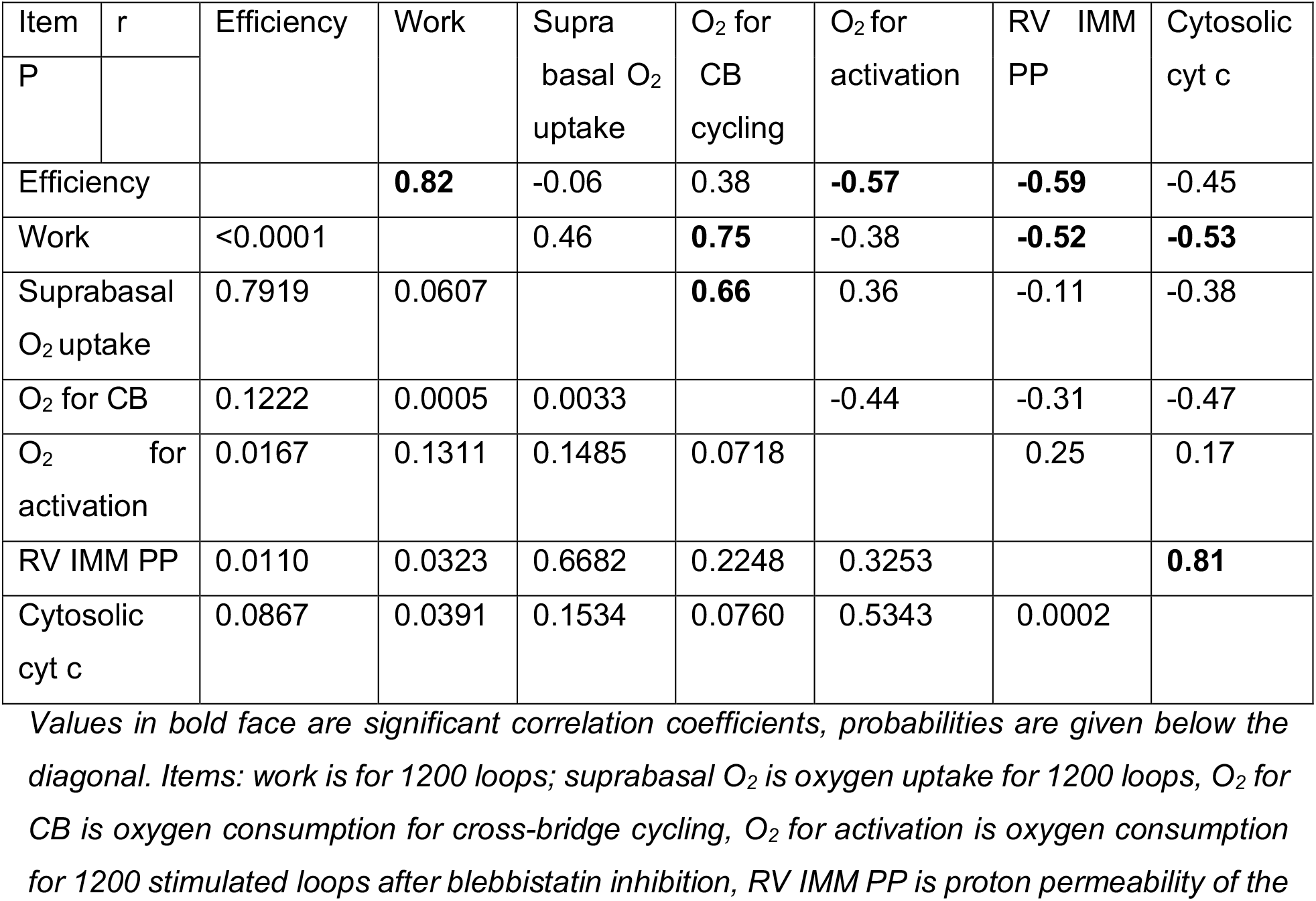
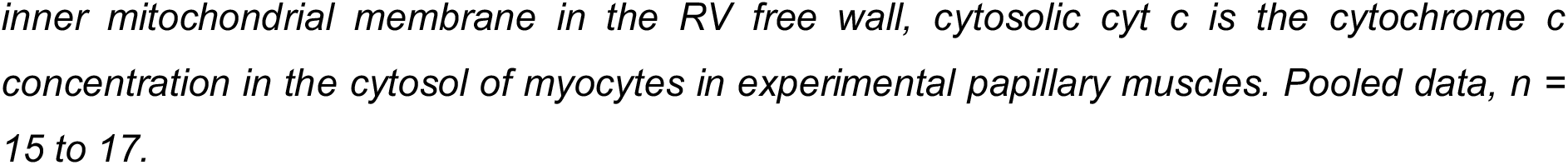
Correlation matrix of variables related to papillary muscle efficiency.

Work was proportional to oxygen consumption for cross-bridge cycling (r^2^ = 0.56). Cytosolic cytochrome c concentration (r^2^ = 0.28) and IMM PP (r^2^ = 0.27) were determinants of work, in agreement with the hypothesis that mitochondrial dysfunction contributes to reduced efficiency. The proportion of variance of work explained by the three significant variables is R^2^ = 0.70 (F_3,13_ = 16.52, p = 0.0051). Significant Deming regressions are shown in Fig. 6.

**Figure 6.**
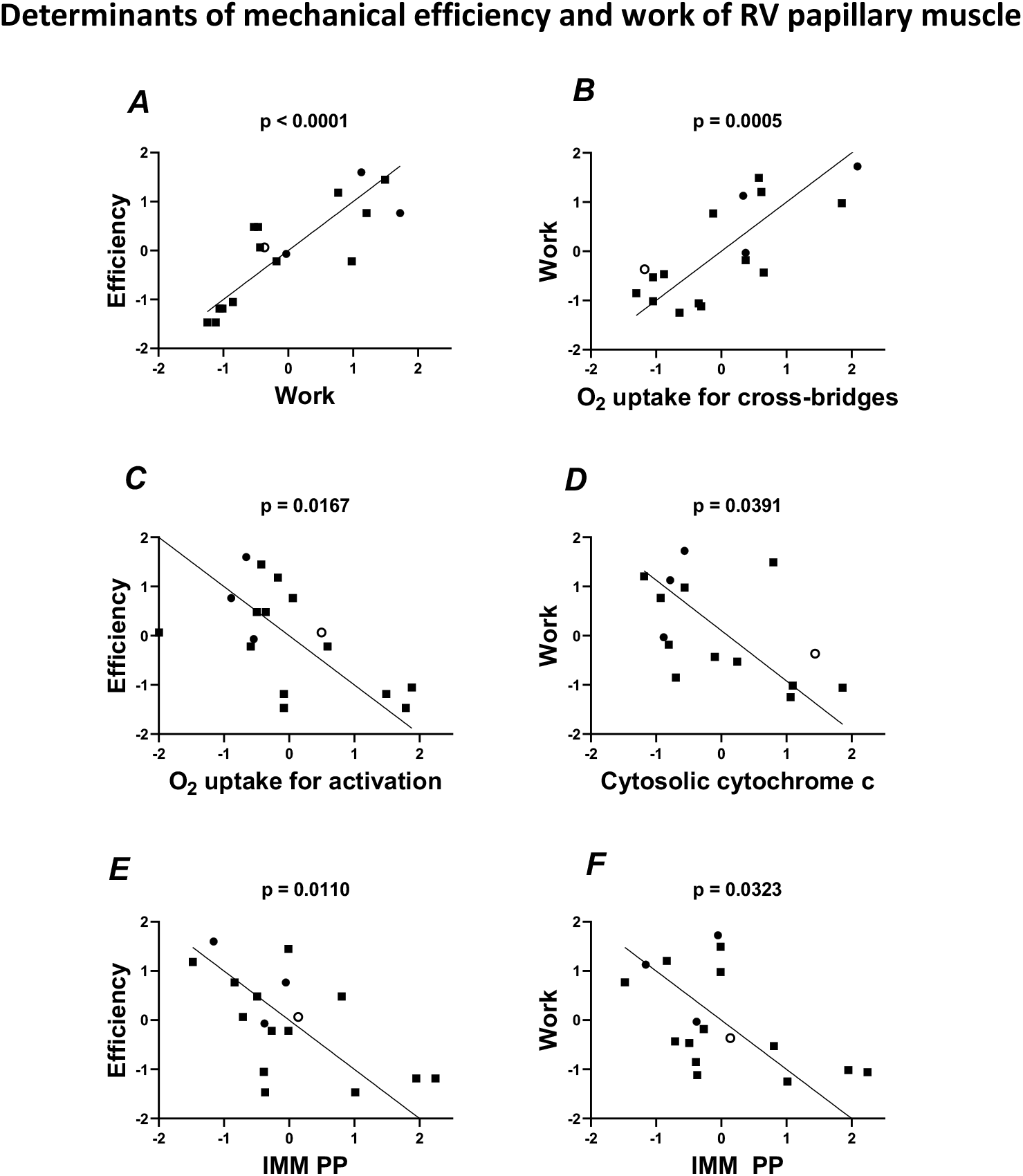
Significant Deming regressions of efficiency versus A: Work, C: oxygen uptake for activation, E: IMM PP (inner mitochondrial membrane proton permeability), and determinants of work: B Oxygen uptake for cross-bridges, D: cytosolic cytochrome c, F: IMM PP. Symbols as in Fig. 2. Data for blebbistatin experiments only.

Because efficiency is calculated from the ratio of work and total oxygen uptake, we also investigated these variances. Initially, the variables also included RV G-6-PDH activity, RV MAO-A activity, RV F_1_F_o_ATPase activity, RV phosphatidylglycerol/cardiolipin, and succinate dehydrogenase (SDH) activity in the experimental muscles. These variables did not correlate with either efficiency, work or total oxygen consumption (P > 0.15).

### Mitochondrial dysfunction and calcium cycling

Cardiolipin metabolism is known to be affected as a result of increased oxidative stress (Paradies et al. 2002). Uncharged reactive oxygen species, H_2_O_2_ and the hydroperoxyl radical HO · (0.3% of the superoxide formed, pK_a_ = 4.8), can permeate the inner mitochondrial membrane and react with double bonds in the acyl chains of cardiolipin (de Grey et al. 2002). MAO-A activity produces H_2_O_2_ on the outer mitochondrial membrane. RV MAO-A activity (for images, see van Eif et al. 2014) was increased in MCT-PH (Table 1) and correlated positively with RV PG/CL (r^2^ = 0.43, P = 0.0141; data in Fig. 1). RV PG/CL correlated strongly with cytosolic cytochrome c in the muscle (Fig. 5, r^2^ = 0.73, P = 0.0015), but was non-linearly related to RV IMM PP (Peters et al. 2019).

To test the possibility that oxygen consumption after blebbistatin inhibition in MCT-PH muscles is due to mitochondrial calcium cycling, oxygen uptake in the presence of ruthenium red was determined. We first studied the effect of ruthenium red on contractility of muscles not treated with blebbistatin in separate experiments in an experimental chamber allowing superfusion of the muscle with Tyrode’s solution with 95% O_2_ and 5% CO_2_ at 37°C. After stabilisation of force at 0.5 Hz stimulation frequency in normal Tyrode’s solution, Tyrode’s solution containing 50 µM ruthenium red decreased force to 16 ± 7 % of initial with 50% reduction after 7.1 ± 3.8 min and 75% reduction after 17 ± 9 min (Fig. 7*A*), in agreement with the observations that ruthenium red also reduced the cytosolic calcium concentration (Tanaka et al. 1997; Malécot et al. 1998). Force recovered irregularly to 49 ± 19% after 1 to 2 hours during washout of ruthenium red, and the colour of the muscle normalized except in damaged myocytes, which remained red. On the basis of these results the muscles in the oxygen chamber were pre-treated for 30 min before oxygen consumption was measured in the presence of ruthenium red. The effect of ruthenium red on oxygen consumption during 4 min of work loops after blebbistatin inhibition is significant (P = 0.0254). It could explain increased blebbistatin-resistant oxygen consumption in two preparations, but not in two others. The effect was small in two preparations which did not recover. As expected on the basis of the results in Fig. 7*A*, recovery of the rate of oxygen consumption after 30 min washout was incomplete, but oxygen uptake after washout and those in the first blebbistatin run were not significantly different (Fig. 7*B*). These results indicate that mitochondrial calcium cycling cannot explain the increase in stimulation-related oxygen uptake in all preparations.

**Figure 7.**
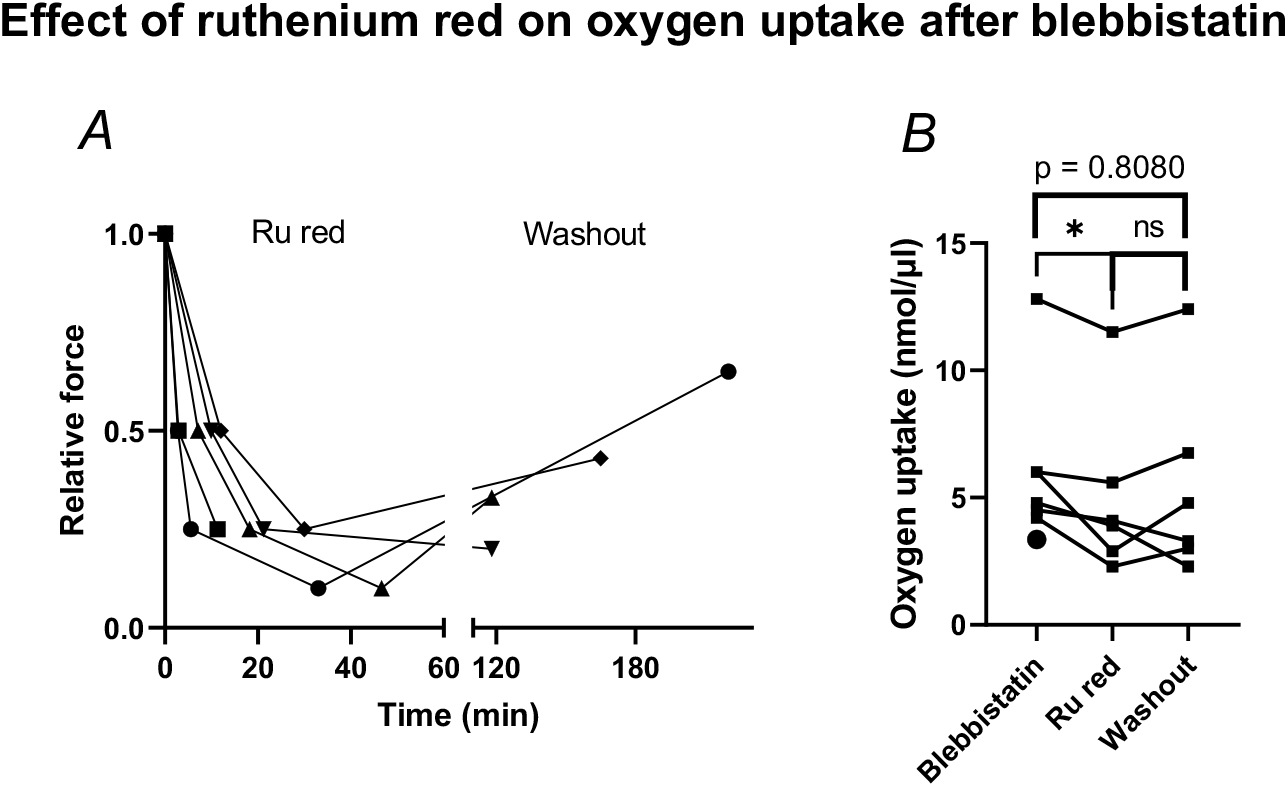
A: Relative force of MCT-PH papillary muscles decreases during 0.5 Hz stimulation in the presence of 50 µM ruthenium red. B: Stimulation-related oxygen uptake for 1200 loops by MCT-PH papillary muscles (after blebbistatin inhibition, see text) is reduced in the presence of ruthenium red and recovered in 4/6 muscles after 30 min washout (paired Students t-tests, *P = 0.0254, ns: P = 0.5147).The circle indicates the mean blebbistatin-resistant oxygen uptake in controls (Table 2).

## DISCUSSION

The results demonstrate that the reduction of myocardial mechanical efficiency with the degree of hypertrophy in MCT-PH is multifactorial, including myocyte cross-sectional area, work, increased energy use for activation, and mitochondrial dysfunction. These factors explain the variance of efficiency statistically. The hypothesis that efficiency is reduced due to increased cytosolic cytochrome c concentration is rejected. The highest cytosolic cytochrome c concentration observed in the muscles was about 50% of the theoretical maximum (for discussion, see van Beek-Harmsen and van der Laarse, 2005). The higher values in the papillary muscle compared to the myocytes in the RV myocardium can partly be explained by the reduction in muscle volume during the experiment (16%, see Methods), but we do not exclude that more cytochrome c has been released during the *in vitro* experiment. However, mechanical efficiency related to cytosolic cytochrome c concentration in myocytes of the RV free wall results in the same conclusion: there is no correlation (r^2^ = 0.01, P = 0.6795, n = 19).

The release of cytochrome c is closely related to the increase of IMM PP in the RV free wall, which can both be due to cardiolipin damage by uncharged reactive oxygen species (Paradies et al., 2002). Prevention of cardiolipin damage may be a therapeutic target. This requires further study.

Considerable variation exists for efficiency within the control and MCT-PH groups, however, oxygen uptake for cross-bridge cycling was relatively constant (Fig. 6*B*). Cross-bridge efficiency calculated from the untransformed data (assuming all experimental error is in the determination of work) was 20% (standard error 2.2%, F_1,16_ = 80.90, P < 0.0001; regression line forced through the origin). This is an underestimate because it includes non-phosphorylating oxygen consumption for mitochondrial proton and calcium cycling and reactive oxygen formation (see below).

Myocyte cross-sectional area varied from 170 µm^2^ in control to 620 µm^2^ in MCT-PH and shows an optimum efficiency at a myocyte cross-sectional area of 420 µm^2^ (at L_opt_), which occurs in both groups. The increase of efficiency with myocyte cross-sectional area in control rats may be related to a decrease of the myocyte surface area to volume ratio (Gibbs et al. 1990). The benificial size effect is reversed with the degree of overload-induced hypertrophy, indicating that the other factors become dominant.

One control muscle had high cytosolic cytochrome c concentration (210 µM). The right-sided myocardium of this rat also had high cytosolic cytochrome c (163 µM), G-6-PDH activity (3.1 µM/s) and IMM PP (32%), which were the highest among control values. This rat stopped breathing before the thorax was opened. The high values could have been due to mild reoxygenation injury. There was no reason to exclude the muscle on the basis of contractile characteristics during the force-length relationship, possibly because mitochondrial capacity was still sufficient.

### Determinants of work

The variation of work was the major determinant of the variance of efficiency. More work increases oxygen consumption for cross-bridge cycling (Fig. 6*B*), reducing the effect on efficiency of increased oxygen uptake for activation (Fig. 6*C*). Work is not related to the cross-sectional area or the interstitial space of the muscle preparation, excluding oxygen diffusion limitation in the muscle as the major factor (van der Laarse et al. 2005; Wong et al. 2010; Han et al. 2011). Hypoxia is also excluded because isoprenaline increased oxygen consumption, and because the rate of oxygen consumption was fairly constant while the oxygen tension in the chamber decreased (Fig. 2*G-I*). At least five alternative explanations for the variation of work exist:

Firstly, cross-bridge activation varies with the magnitude of calcium transients. Calcium transients are increased in MCT-PH (Power et al. 2018; Sabourin et al. 2018), partly explaining the increased oxygen consumption after blebbistatin inhibition. However, variation of work is also found among control muscles. Secondly, cross-bridge activation varies with myofibrillar calcium sensitivity. Calcium sensitivity can increase with adrenergic stimulation, but can decrease with increasing heart rate in MCT-PH (Lamberts et al. 2007). Thirdly, ATP delivery to myosin ATPase is reduced. This would reduce the rate of cross-bridge detachment, shortening velocity and power output. Mitochondrial function is impaired, as indicated by increased cytosolic cytochrome c and IMM PP and the phosphocreatine shuttle malfunction (Fowler et al. 2015, 2019). It is a possibility that the available energy is used preferentially for increased calcium cycling when facilitated energy transport fails. Furthermore, because glucose uptake during the efficiency determination can be insufficient, due to diffusion limitation in the extracellular space and the lack of insulin in the Tyrode’s solution, the muscles may use glycogen stores during the 5 Hz activation period (Goodwin et al. 1998). The calcium transients liberate glucose 6-phosphate from glycogen by activating phosphorylase kinase A, but this could be less than the amount needed to match energy demand. Therefore, it may be that the production of acetyl CoA is limiting oxidative phosphorylation. Fourthly, a shift of fast myosin type α to slow myosin type β occurs in right-sided myocardium in MCT-PH (Kögler et al. 2003, Sabourin et al. 2018), which can cause a reduction of power output. Fifthly, cross-bridge cycling may be inhibited by post-translational modifications (Lin et al. 1991; McClellan et al. 1996). Interaction of these possiblilties is not excluded.

We exclude experimentally induced hypoxia during the efficiency determination as a cause of reduced work. However, myocyte hypoxia cannot be excluded in hypertrophied RV myocardium *in vivo* (Oknińska et al. 2021). The positive correlation of work and efficiency may suggest that inotropic intervention improves both cardiac output and efficiency. In case the intervention leads to increased myocardial oxygen demand – e.g. as observed in the isoprenaline experiments – mitochondrial dysfunction may be accelerated.

### Comparison of oxygen consumption and heat production for activation

Although differences between the present experiments and the heat experiments of Pham et al. (2017, 2018) exist, a comparison of activation heat and blebbistatin-resistant oxygen consumption reported in the present study is possible, assuming that calcium transients are similar and the energy required for activation processes is produced by the oxidation of glucose (no mechanical work produced, cf. Woledge et al. 1985, p. 243). [The experiments in Auckland and Amsterdam were carried out independently. Both labs induced pulmonary hypertension by 60 mg/kg MCT in Wistar rats. Body masses and time between MCT injection and experiment differed. We used N-acetyl cysteine in the Tyrode solution and a different dissection and mounting procedure. Preparation and shortening protocols differed, but stimulus frequency was 5 Hz, Ca^2+^ in the Tyrode solution was 1.5 mM, glucose 10 mM, pH = 7.4 and experimental temperature was 37°C. Volume determinations of the preparations were similar.]

Activation heat per stimulus after blebbistatin inhibition in control trabeculae was 1.2 ± 0.7 kJ/m^3^ (n = 4, Fig. 4*D*, Pham et al. 2017) or 1.0 ± 0.6 kJ/m^3^ (n = 14, Fig. 3*F*, Pham et al. 2018). The weigthed mean ± SD calculated from these data is 1.1 ± 0.6 kJ/m^3^. When the control muscle with high cytosolic cytochrome c (and G-6PDH activity and IMM PP in the RV wall) is excluded, we find 2.8 ± 0.6 mmol/m^3^ oxygen uptake per stimulus after blebbistatin inhibition in three papillary muscles (Table 2). Thus 1.1/2.8 10^−3^ = 393 kJ/mol O_2_ would be produced, approximating (83 %) the expected heat of combustion of glucose (473 kJ/mol O_2_). The ratio of activation heat in preparations of MCT-PH rats (1.5 ± 0.6 kJ/m^3^, n = 21, Fig. 3*F* in Pham et al. 2018) and blebbistatin-resistant oxygen consumption (5.8 ± 3.9 mmol/m^3^, Table 2) equals 259 kJ/mol O_2_, 55% of the heat of combustion of glucose. The discrepancy between calculated and expected heat/O_2_ suggests that oxygen consumed for activation processes in compensated and failing heart muscle cannot all be used to oxidise glucose to CO_2_ and H_2_O. In an extreme case, all glucose is oxidised to CO_2_ and H_2_O_2_, which requires 9 O_2_/glucose and produces 191 kJ/mol O_2_ (according to Hess’s law and enthalpy of formation of H_2_O_2_ = - 187kJ/mol; ATcT, 2021). This can occur when electrons cannot be transported to complex IV due to loss of cytochrome c and react with O_2_ at complexes I, II and/or III (Zhao et al., 2003). This explanation of the value of activation heat/O_2_ in MCT-PH myocardial preparations requires that any superoxide formed is converted to H_2_O_2_ and that H_2_O_2_ diffuses into the Tyrode solution. H_2_O_2_ diffusing into coronary and myocardial lymph vessels may be clinically relevant, e.g. it may increase glutathione disulphide (GSSG) in erythrocytes, and interfere with nitric oxide signaling.

Reactive oxygen species can also upregulate G-6-PDH activity by oxidative deactivation of glyceraldehyde 3-phosphate dehydrogenase (Eaton et al. 2002), thereby stimulating the production of NADPH in the pentose phosphate pathway (Ralser et al. 2007). The production of reactive oxygen species in excess of the capacity of scavengers (suggested by the G-6-PDH activity in Table 1) can explain the progression of mitochondrial dysfunction by reacting with cardiolipin’s double bonds. Once cytochrome c is released from the mitochondria, the process is potentially autocatalytic.

The comparison of heat production and oxygen consumption for activation suggests that two-thirds of the suprabasal glucose consumed by the overloaded right-sided papillary muscle is converted to CO_2_ and H_2_O_2_, which can exceed the scavenging capacity of the glutathione antioxidant system. This also suggests that using the heat of combustion of glucose (or the commonly used 20 J/ml O_2_) as a conversion factor may underestimate mechanical efficiency of failing hearts. In view of the variance within experimental groups, future examination of these possibilities requires paired observations of work, heat and hydrogen peroxide release as well as oxygen consumption.

## Additional information

The efficiency experiments were carried out at the Department of Physiology, VU University Medical Center, Amsterdam

## Acknowledgements

We thank F.D. Dijkema and P.M. Sneekes for expert technical assistance.

## Author contributions

Conception and design of the study WJvdL. IS and AVN provided and characterised experimental rats. WJvdL and SJPB carried out efficiency experiments and histochemistry. FMV performed HPLC-MS experiments. DvG designed hardware and wrote software. WJvdL, SJP, IS, FMV and DvG analysed data. WJvdL wrote the paper. All authors critically reviewed the manucript and agree with the final version.

